# Local aromatic interactions define temperature sensitivity of phase separation in an intrinsically disordered protein

**DOI:** 10.64898/2026.05.07.723405

**Authors:** Yumiko Ohhashi, Suguru Nishinami, Yoko Maruyama, Mao Fukuyama, Kentaro Shiraki, Eri Chatani, Hideki Taguchi

## Abstract

Liquid–liquid phase separation (LLPS) of intrinsically disordered proteins is highly sensitive to environmental conditions, yet the molecular basis of sharp temperature responsiveness remains poorly understood. Here, we investigate a sequence-encoded mechanism underlying the pronounced temperature sensitivity of phase separation using Sup35NM, the intrinsically disordered domain of the yeast prion Sup35. We show that a tyrosine-rich local structural region within this domain encodes strong temperature responsiveness of droplet formation. Mutational analyses reveal that tyrosine residues mediate both intramolecular interactions that stabilize local structure and intermolecular interactions required for LLPS. Substitution of the tyrosine residues with alanine disrupts local structure and weakens intermolecular interactions, thereby diminishing temperature sensitivity. In contrast, substitution with phenylalanine promotes rapid droplet gelation, abolishes internal fluidity, suppresses amyloid formation, and confers resistance to temperature-induced dissolution. Based on these findings, we propose a molecular model in which finely tuned, moderately weak aromatic interactions among tyrosines enable reversible local compaction that is sensitive to temperature, thereby generating a sharp phase transition. These results suggest that amino acid sequences encode not only phase separation propensity but also the sensitivity of condensates to environmental perturbations, providing a framework for understanding how intrinsically disordered proteins act as molecular sensors of cellular conditions.

## INTRODUCTION

The crowded intracellular environment makes it difficult for diffusing proteins, nucleic acids, and other macromolecules to participate in essential biological reactions. To overcome this limitation, cells utilize liquid-liquid phase separation (LLPS) to form membraneless compartments that partition in the cytoplasm or nucleus, locally concentrating the molecules required for a series of biochemical processes. These phase-separated droplets act as molecular reservoirs, organizational centers, and reaction crucibles, enabling the smooth progression of cellular processes (*1–4*). Many key proteins involved in intracellular droplet formation contain low-complexity domains (LCDs), which are composed of a limited set of amino acids. These LCDs sustain droplet state through transient and multivalent interactions (*5–7*). In vitro, proteins containing LCDs frequently undergo phase separation into a dense phase, where proteins are highly concentrated within droplets, and a surrounding dilute phase. Amyloid-like aggregates have been observed to form within the dense phase of several proteins (*4,8–11*). Since droplets with specific proteins have been implicated as potential precursors of pathological aggregates associated with neurodegenerative diseases, cells are thought to possess regulatory mechanisms that prevent harmful aggregation originating from droplets. One such mechanism could be the environmentally responsive regulation of the droplet formation and dissolution, preventing unnecessary persistence of intracellular droplets.

The formation and dissolution of intracellular droplets are tightly regulated by environmental cues and cellular reactions (*3,4*). Various factors contribute to this regulation, including alterations in protein solubility induced by post-translational modifications, the addition or removal of intermolecular interaction sites (*12–14*), modulation by other proteins, such as molecular chaperones (*12,15*), and environmental stimuli, such as pH and ionic strength (*16,17*). In vitro phase separation is highly sensitive to environmental conditions, including ionic strength, pH, pressure, and temperature (*18,19*). For example, electrostatic interactions play an important role in droplet formation for many proteins; therefore, ionic strength and pH are critical environmental factors (*20–23*). Temperature dependence is another hallmark of phase separation. In the experimental temperature range, many proteins form droplets at low temperatures, whereas others do so at elevated temperatures. These temperature-dependent behaviors are determined by the intermolecular interactions that drive droplet formation, which are encoded in the amino acid sequence (*24–27*). Temperature-responsive phase separation is a common property of many proteins, characterized by a sharp transition in droplet volume over a narrow temperature range. However, the molecular mechanism underlying this pronounced temperature-sensitivity remains poorly understood.

The yeast prion protein Sup35 provides a useful model system to address this question. Sup35 from *Saccharomyces cerevisiae* forms phase-separated droplets in cells under stress conditions such as carbon starvation, low pH, and high osmotic pressure (*17,28,29*). Sup35 consists of three domains: N, M, and C. The N and M domains (Sup35NM) are intrinsically disordered regions responsible for both phase separation and amyloid formation. Within the N domain, a compact local structure has been identified that serves as a site for intermolecular interactions during oligomerization (*30,31*). Furthermore, the N domain forms the core of intermolecular interactions during amyloid fibril formation (*30,32*), and is therefore referred to as the prion domain. This domain is typical LCD, highly enriched in glutamine, glycine, asparagine, and tyrosine residues, which together account for approximately 80% of the sequence. Similar compositional features are found in several human RNA-binding proteins implicated in neurodegenerative diseases, such as FUS and hnRNPA1 (*33,34*), suggesting that common physicochemical principles may underlie phase behavior across diverse IDPs. The M domain is characterized by a high density of charged residues, with a charge gradient: a predominance of positive charge at the N-terminal side and negative charge at the C-terminal side. This charge distribution could facilitate intermolecular electrostatic interactions. In vitro studies of Sup35NM have demonstrated that weakly acidic conditions favor droplet formation (*17*). Our previous studies revealed that Sup35NM spontaneously forms amyloid fibrils within droplets several hours after droplet formation (*35,36*).

In addition, droplet formation of Sup35NM is enhanced at low temperatures and under polyethylene glycol (PEG) -induced crowding conditions.

In this study, we investigated the molecular mechanism underlying the pronounced temperature-sensitivity of Sup35NM droplet formation. Our findings demonstrate that tyrosine residues within the local structure (LS) region of the N domain play a crucial role in this temperature sensitivity. These tyrosine residues participate in both intra- and intermolecular aromatic interactions, contributing significantly to droplet formation. Weakening tyrosine-mediated interactions disrupted intermolecular interactions among N-domains, shifting Sup35NM droplet formation to M-domain–driven and attenuating temperature sensitivity, whereas strengthening these interactions induced rapid gelation, rendering droplets temperature-resistant. Based on these observations, we propose that moderately weak, temperature-responsive interactions between tyrosine residues are essential for the pronounced temperature sensitivity of Sup35NM phase separation.

## RESULTS

### The local structure region within the N domain confers strong temperature sensitivity on droplet formation

As shown in our previous study, Sup35NM-mediated droplet formation was monitored as an increase in turbidity (*36*).Temperature-dependent turbidity measurements were performed by increasing the temperature in 5°C increments from 20°C to 60°C. Optical microscopy confirmed that droplet abundance decreased concomitantly with turbidity (Fig. 1, A and B). We found that, under 15% PEG conditions, Sup35NM droplets formed at low temperatures rapidly disappeared between 40°C and 50°C (Fig. 1, A and B). This sharp decline within such a narrow temperature range prompted us to investigate the underlying mechanism. We hypothesized that this strong temperature-sensitivity is encoded in the amino acid sequence of Sup35NM. To identify the responsible regions, we examined the droplet formation by the N and M domains individually (Fig. 1, A and B). Both domains were capable of undergoing phase separation independently under our experimental conditions. The M domain exhibited PEG concentration dependence similar to that of Sup35NM, whereas such dependence was not observed for the N domain (fig. S1). Under 15% PEG conditions, the N domain exhibited a sharp decrease in droplet abundance between 20°C and 30°C. In contrast, the M domain showed gradual and monotonic temperature dependence, and droplets remained even at 60°C (Fig. 1B), a temperature at which droplets formed by Sup35NM and the N domain had completely dissolved (Fig. 1B). These findings suggest that the N domain confers the temperature-sensitivity to the Sup35NM phase-separation.

**Figure 1.**
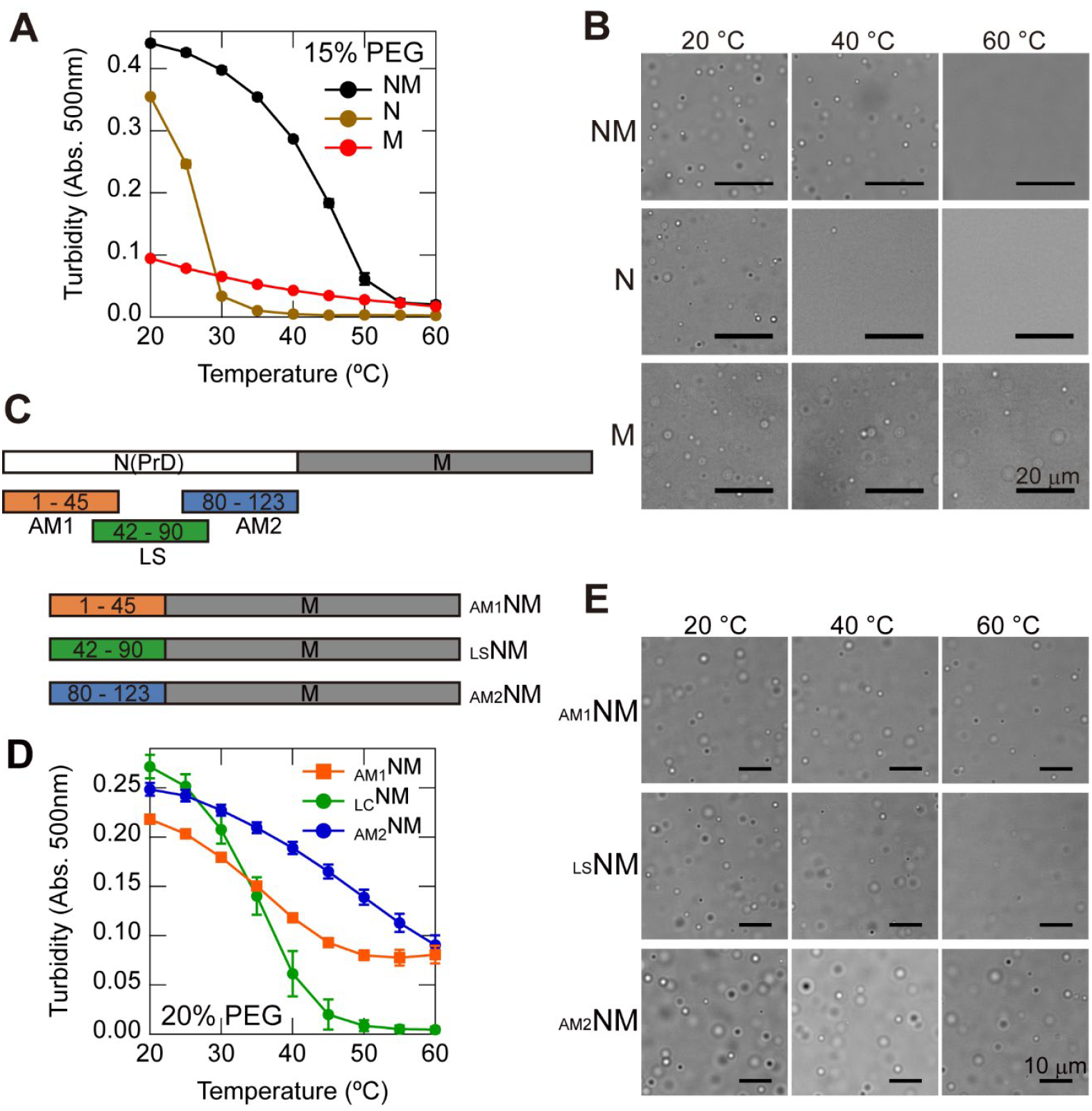
The LS region of the N domain confers strong temperature-sensitivity to Sup35NM droplet formation. (**A**) Temperature-dependent droplet formation of Sup35 NM, N, and M domains monitored by turbidity at 500 nm in the presence of 15% (w/v) PEG 20,000. (**B**) Representative optical microscopy images showing temperature-dependent changes in droplet abundance for Sup35NM, N, and M domains under 15% (w/v) PEG 20,000 conditions. Scale bars, 20 µm. (**C**) Schematic illustration of truncated Sup35NM variants: AM1NM, LSNM, and AM2NM. (**D**) Temperature-dependent turbidity profiles of AM1NM, LSNM, and AM2NM measured in 20% (w/v) PEG 20,000. (**E**) Representative optical microscopy images showing temperature-dependent changes in droplet abundance for AM1NM, LSNM, and AM2NM under 20% (w/v) PEG 20,000 conditions. Scale bars, 10 µm.

To determine which region of the N domain responsible for the temperature sensitivity, we divided the N domain into three regions, residues 1-45, 42-90, and 80-123, and fused each segment to the M domain to generate three deletion mutants (Fig. 1C). The 1-45 and 80-123 regions, known to be involved in intermolecular interactions during amyloid fibril formation (*30, 37*), were designated AM1 and AM2, respectively. The 42-90 region, shown to form a compact local structure (*30, 31*), was designated LS. The resulting fusion proteins were termed _AM1_NM, _AM2_NM, and _LS_NM (Fig. 1C). Turbidity measurements revealed that _LS_NM exhibited temperature-sensitivity comparable to that of Sup35NM, with a sharp decrease in turbidity between 30°C and 40°C under 20% PEG conditions (Fig. 1, D and E). In contrast, _AM1_NM and _AM2_NM exhibited monotonic temperature-dependent turbidity changes similar to those of the M domain, and droplets remained even at 60°C (Fig. 1, D and E). Together, these results suggest that the LS region of the N domain plays a central role in conferring strong temperature-sensitivity on Sup35NM droplet formation.

### Tyrosine residues in the LS region mediate temperature sensitivity and molecular compactness of Sup35NM

To elucidate the mechanism underlying the strong temperature-sensitivity conferred by the LS region, we examined the effects of amino acid substitutions. The Sup35 N domain is a typical LCD enriched in Gln, Gly, Asn, and Tyr. Notably, the LS region contains 12 Tyr residues, substantially more than AM1 (6 residues) or AM2 (5 residues) regions (fig. S2). To evaluate the contribution of Tyr in the LS region to temperature sensitivity, we generated mutants in which all Tyr residues were substituted with Ala (YA-NM) or in which half were substituted (YAh-NM) (Fig. 2A). Turbidity measurements showed that both Ala mutants displayed gradual and monotonic temperature dependence, indicating reduced temperature-sensitivity (Fig. 2B). When these substitutions were introduced into the isolated N domain (Fig. 2C), turbidity increases were completely abolished, indicating that substitution of all, or even half, of the Tyr to Ala disrupts intermolecular interactions among N domains (Fig. 2D). These results suggest that YA-NM and YAh-NM undergo droplet formation primarily driven by the M domain, without inter N domain interactions, thereby resulting in diminished temperature sensitivity.

**Figure 2.**
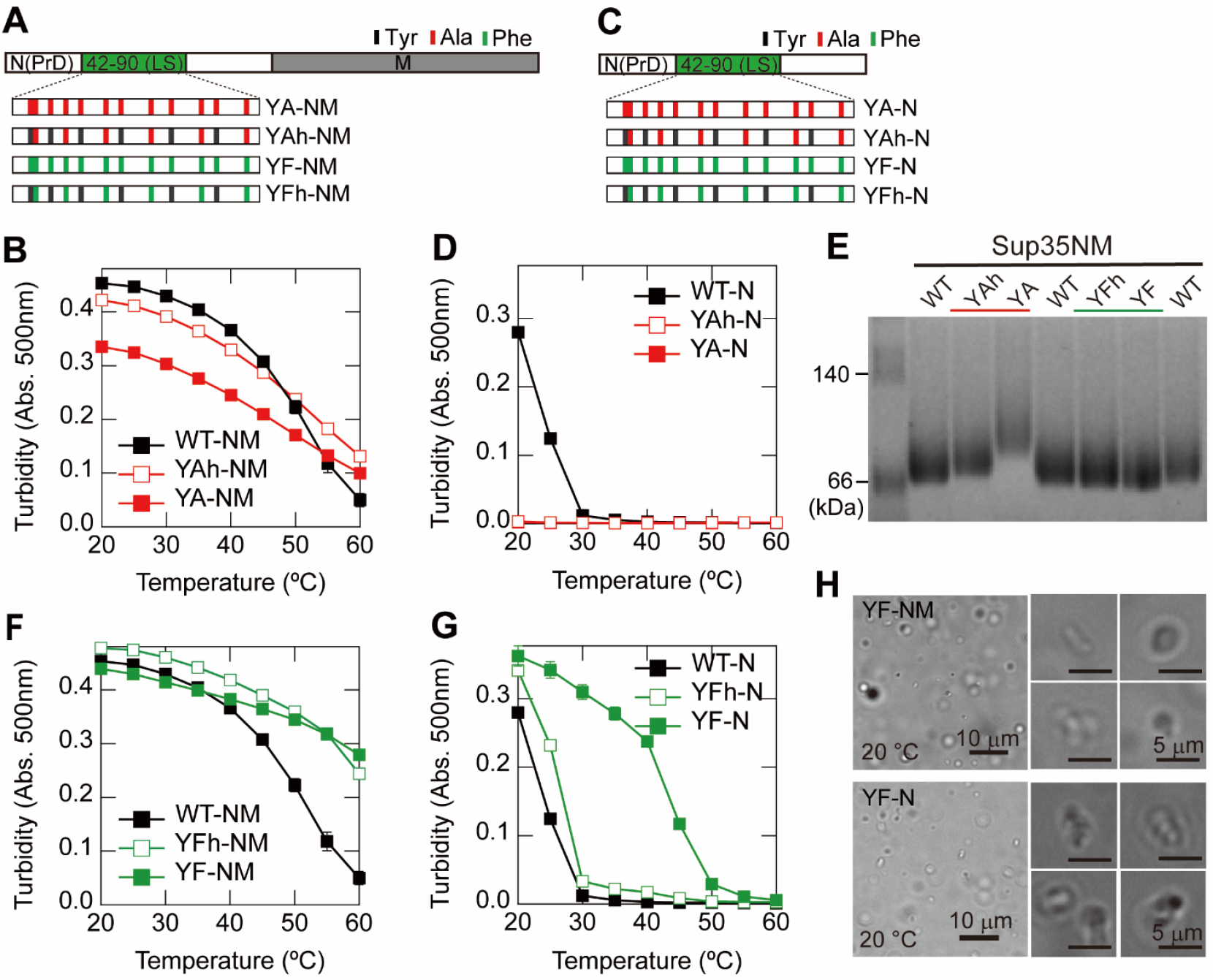
Tyrosine substitutions in the LS region alter droplet properties of Sup35. (**A**) Schematic representation of Tyr-substituted Sup35NM mutants (YA-NM, YAh-NM, YF-NM, and YFh-NM). Tyr positions are indicated by black lines, Ala substitutions by red lines, and Phe substitutions by green lines. (**B**) Temperature-dependent turbidity profiles of Tyr-to-Ala Sup35NM mutants (YA-NM and YAh-NM) in 20% (w/v) PEG 20,000. (**C**) Schematic representation of Tyr-substituted Sup35N mutants (YA-N, YAh-N, YF-N, and YFh-N). (**D**) Temperature-dependent turbidity profiles of Tyr-to-Ala Sup35N mutants, YA-N and YAh-N in 20% (w/v) PEG 20,000. (**E**) Blue-Native PAGE analysis of Sup35NM mutants carrying Tyr-to-Ala (red) or Tyr-to-Phe (green) substitutions. (**F**) Temperature-dependent turbidity profiles of Tyr-to-Phe Sup35NM mutants (YF-NM and YFh-NM) in 20% (w/v) PEG 20,000. (**G**) Temperature-dependent turbidity profiles of Tyr-to-Phe Sup35N mutants (YF-N and YFh-N) in 20% (w/v) PEG 20,000. (**H**) (left) Representative optical microscopy images showing droplet morphology of YF-NM and YF-N mutants at 20°C in 20% (w/v) PEG 20,000. Scale bars, 10 µm. Right panels show magnified views. Scale bars, 5 μm.

To examine whether Tyr-to-Ala substitutions in the LS region induce conformational change, we analyzed the proteins using blue-native polyacrylamide gel electrophoresis (Blue-Native PAGE) (Fig. 2E). WT-Sup35NM (WT-NM), with a calculated molecular weight of 30 kDa, exhibited a pronounced band shift to an apparent molecular weight of approximately 66 kDa, likely reflecting the extended conformation of regions outside the LS region, which are predominantly disordered. The YA-NM mutant displayed an even greater shift toward higher apparent molecular weights, and the YAh-NM mutant showed an intermediate shift. These observations suggest that Tyr-to-Ala substitutions disrupt local structural compaction, leading to a more expanded molecular conformation. Circular dichroism spectroscopy (fig. S3) at 20°C and 60°C showed that both WT-NM and YA-NM exhibited spectra characteristic of random coil conformations, with no appreciable differences between them. This finding indicates that the N domain does not adopt a defined secondary structure but instead maintains a compact conformation through intramolecular interactions. Taken together, these findings suggest that the compactness of the LS region arises from intramolecular interactions among Tyr aromatic rings, which are mechanistically analogous to the intermolecular interactions that drive phase separation.

### Tyr to Phe substitution enhances droplet gelation via strengthened intermolecular interactions

To further investigate how Tyr-mediated interactions contribute to temperature sensitivity, we analyzed mutants in which all Tyr residues in the LS region were substituted with Phe (YF-NM) as well as mutants in which half were replaced (YFh-NM) (Fig. 2A). While Phe can participate in aromatic interactions similar to Tyr, it is more hydrophobic, interactions among Phe residues are generally stronger than among Tyr (*38, 39*). Turbidity measurements revealed that both YF-NM and YFh-NM exhibited modest temperature-dependent changes in turbidity, indicating reduced temperature-sensitivity (Fig. 2F). Blue-Native PAGE analysis showed that YF-NM migrated slightly toward lower apparent molecular weight compared to WT-NM, suggesting that the LS region adopts a more compact conformation (Fig. 2E). To further assess the effects of these substitutions, we generated isolated N-domain mutants (YF-N and YFh-N) (Fig. 2C). YF-N displayed a gradual decrease in turbidity at lower temperatures, followed by a sharp decline near 40°C, and turbidity became undetectable at 60°C. This result suggests that Tyr-to-Phe substitution increases the thermal stability of droplets, resulting in an elevated temperature threshold for droplet dissolution. YFh-N exhibited a turbidity profile similar to that of WT-N, although a small subset of droplets displayed behavior resembling that observed for YF-N (Fig. 2G). Optical microscopy revealed that droplets formed by YF-NM and YF-N deviated from the typical spherical morphology and appeared distorted (Fig. 2H), suggesting reduced internal fluidity and transition toward a gel-like state.

The process of gelation, characterized by the formation of solid or semisolid assemblies, has been observed during droplet maturation in various proteins and is often associated with the emergence of amyloid-like cross-β structures (*5, 40, 41*). To explore this possibility, we stained droplets with Thioflavin T (ThT), a dye that selectively binds to cross-β structures, and performed fluorescence microscopy observations (Fig. 3A). Thirty minutes after droplet formation, YF-NM droplets exhibited significantly stronger ThT fluorescence than WT-NM droplets, indicating the presence of cross-β structures. To directly assess the internal fluidity of droplets, we performed fluorescence recovery after photobleaching (FRAP) using Alexa Fluor 488-labeled WT-NM, YF-NM, and YA-NM (Fig. 3, B and C). WT-NM droplets exhibited clear fluorescence recovery, consistent with liquid-like behavior. In contrast, YF-NM droplets showed no detectable recovery, indicating a complete loss of internal fluidity. YA-NM droplets exhibited accelerated fluorescence recovery compared to WT-NM, suggesting increased fluidity. Upon prolonged incubation, WT-NM and YA-NM formed amyloid fibrils, whereas YF-NM remained as gel-like aggregates without fibril formation (Fig. 3D). Collectively, these findings suggest that a Tyr-to-Phe substitution in the LS region promotes a rapid gelation upon droplet formation, leading to loss of internal fluidity and reduced temperature sensitivity. The enhanced stability of these gel-like assemblies appears to prevent subsequent structural rearrangements into amyloid fibrils, likely due to strengthened intermolecular interactions mediated by Phe residues.

**Figure 3.**
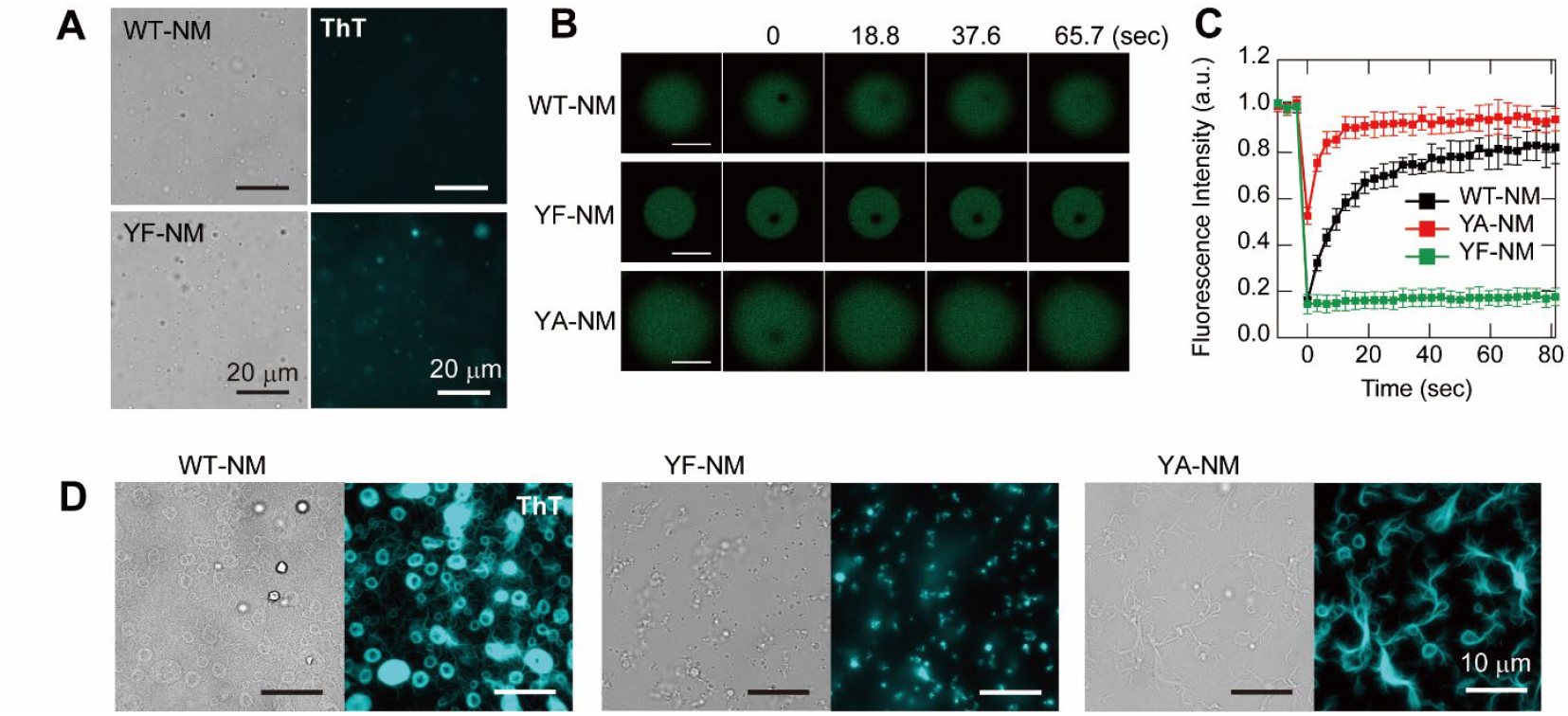
Phenylalanine substitution in the LS region promotes gelation of Sup35NM droplets. (**A**) ThT fluorescence of droplets formed by WT-NM and YF-NM immediately after droplet formation. (**B**) Representative fluorescence images from fluorescence recovery after photobleaching (FRAP) experiments of WT-NM, YF-NM, and YA-NM droplets labeled with Alexa Fluor 488, showing pre-bleach, immediately post-bleach (0 sec), and recovery tike points. Scale bars, 5 μm (**C**) Normalized fluorescence recovery curves obtained from FRAP measurements of WT-NM, YF-NM, and YA-NM droplets. Data are presented as mean ± SD (n = 7). (**D**) Representative ThT Fluorescence images showing droplet morphology of WT-NM, YF-NM, and YA-NM at 2 days after droplet formation. Scale bars, 10 μm.

### Urea weakens intermolecular interactions in YF mutant droplets, restoring temperature-sensitivity

Given that Phe substitutions strengthened intermolecular interactions and promoted gelation, we next examined whether denaturants could reverse this effect. As expected, addition of 2 M urea significantly reduced ThT fluorescence in YF-NM droplets (Fig. 4A), indicating an attenuation of cross-β–like interactions within the droplets. Although droplet abundance was modestly reduced, droplets were still observed under this condition. Moreover, the addition of urea suppressed the gelation of YF-NM droplets, preserving their spherical morphology (Fig. 4A), and induced amyloid formation approximately 1 h after droplet formation (Fig. 4B, fig. S4).

**Figure 4.**
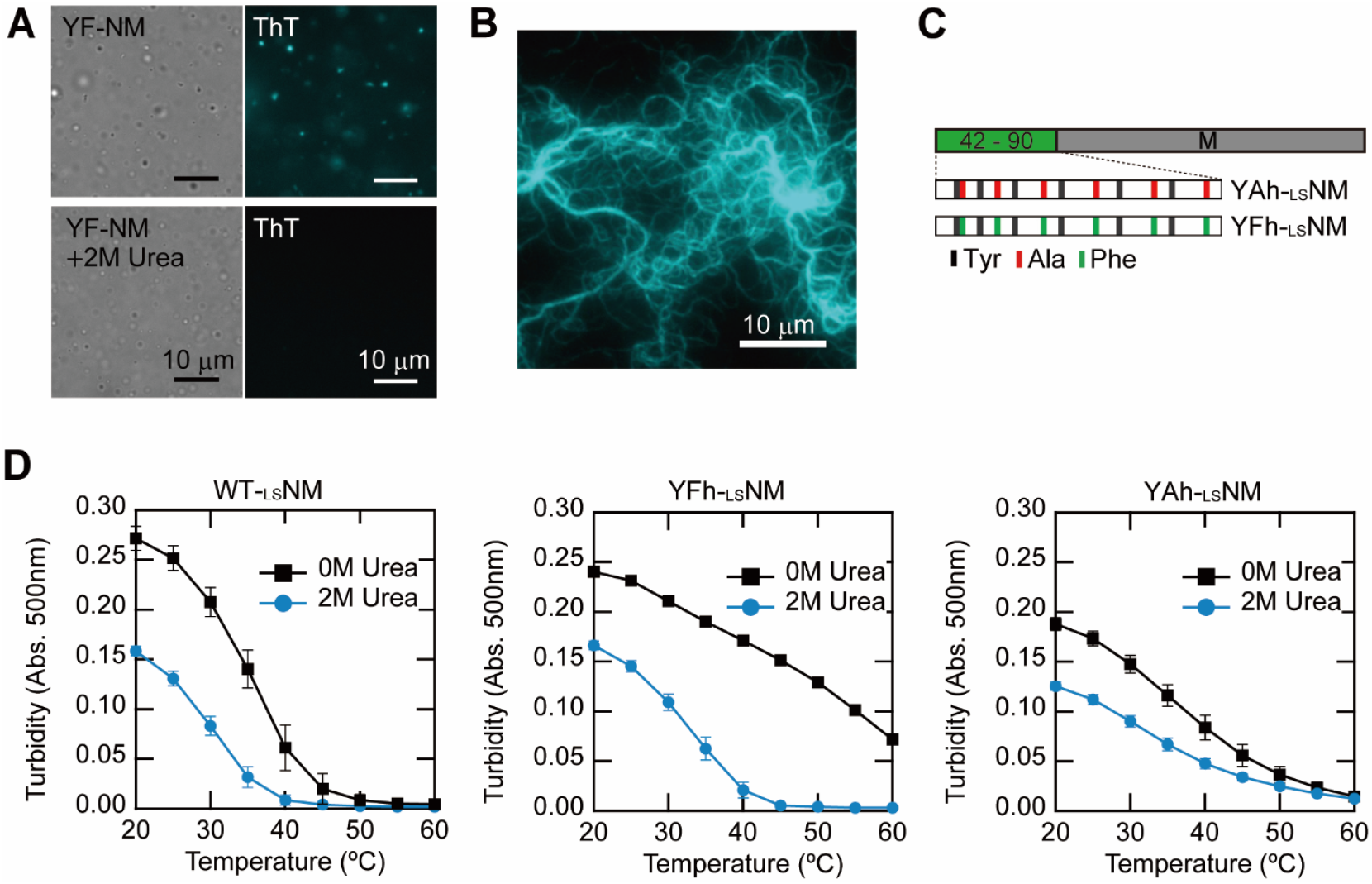
Denaturant restores temperature sensitivity in Phe-substitute mutants. (**A**) ThT fluorescence images of YF-NM droplets formed in 20% (w/v) PEG 20,000 in the absence (top) or presence (bottom) of 2 M urea. Scale bars, 10 μm. (**B**) ThT fluorescence images of amyloid fibrils formed by YF-NM in the presence of 2 M urea. Scale bars, 10 μm. (**C**) Schematic representation of _LS_NM mutants (YAh-_LS_NM and YFh-_LS_NM). (**D**) Temperature-dependent turbidity profiles of WT-NM, YF-NM, and YA-NM in 20% (w/v) PEG 20,000 in the absence (black) or presence (blue) of 2 M urea.

These observations suggest that urea weakens intermolecular interactions among Phe within YF-NM droplets, restores internal fluidity, and permits structural rearrangement toward amyloid formation. Based on these findings, we predicted that, in the presence of urea, the temperature dependence of YF-NM turbidity would resemble that of WT-NM. However, rapid amyloid formation prevented reliable turbidity measurements of YF-NM. We therefore assessed temperature sensitivity using _LS_NM Ala and Phe mutants (YAh-LSNM, YFh-LSNM) (Fig. 4, C and D). Both mutants exhibited monotonic temperature dependence, similar to YA-NM and YF-NM. YFh-_LS_NM droplets displayed detectable ThT fluorescence, YAh-_LS_NM showed disrupted local structure in Blue-Native PAGE (fig. S5). Upon addition of 2 M urea, YFh-_LS_NM acquired strong temperature sensitivity, displaying turbidity behavior comparable to WT-_LS_NM(Fig. 4D). In contrast, YAh-_LS_NM maintained a gradual and monotonic turbidity change even after urea addition. Collectively, these findings demonstrate that the reduction in temperature sensitivity observed in the Phe mutants arises from strengthened aromatic interactions and the consequent rapid gelation, which are reversible upon urea addition.

### Temperature sensitivity cooperates with additional environmental factors to regulate droplet formation

Finally, we investigated how temperature-sensitivity coordinates with other environmental factors to regulate droplet formation. Specifically, we analyzed the effects of pH, denaturant concentration, and ionic strength on droplet formation by Sup35NM and Sup35M using temperature-dependent turbidity assays (Fig. 5, fig. S6). In both proteins, droplet formation was suppressed under basic pH conditions and at elevated concentrations of urea or NaCl, as reflected by downward shifts in the droplet formation temperature (Fig. 5, fig. S6). In the presence of NaCl, Sup35NM, characterized by a strong temperature-sensitivity, retained its sharp temperature dependence while droplet formation temperature shifted to lower values, this shift expanded the temperature range in which droplets were absent (Fig. 5, A and B). In contrast, Sup35M, which exhibits weak temperature-sensitivity, maintained droplet formation over a broad temperature range even in the presence of NaCl, and droplet-free conditions were not observed within the tested condition range (Fig. 5, C and D). Collectively, these results suggest that proteins exhibiting strong temperature sensitivity in LLPS are also more responsive to additional environmental stimuli, enabling tighter regulation of droplet formation across varying conditions.

**Figure 5.**
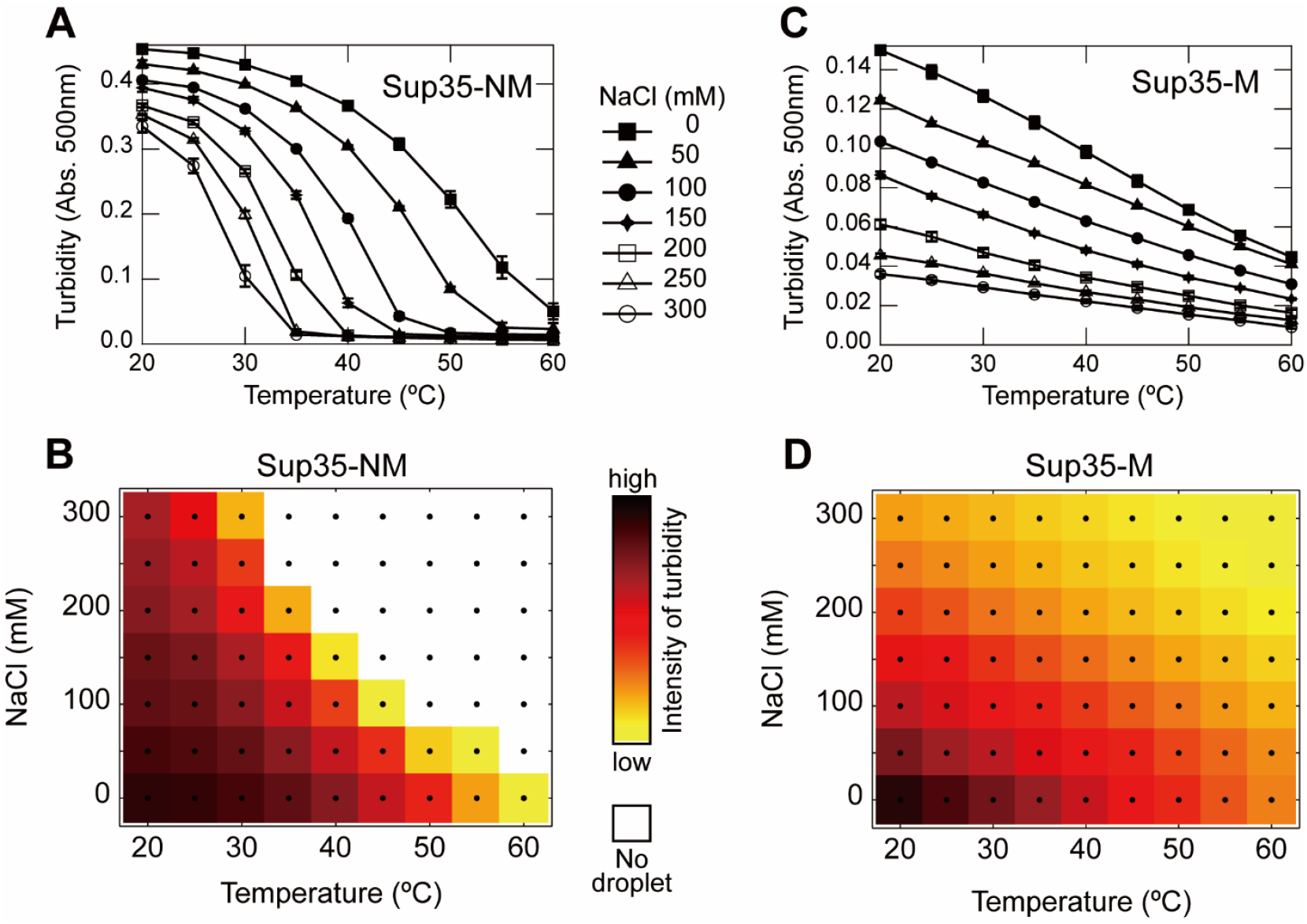
Temperature sensitivity cooperates with additional environmental factors. (**A**) NaCl concentration dependence of Sup35NM droplet formation measured by turbidity. (**B**) Phase diagram of Sup35NM droplet formation constructed from turbidity measurements in (A). (**C**) NaCl concentration dependence of Sup35M droplet formation measured by turbidity. (**D**) Phase diagram of Sup35M droplet formation constructed from turbidity measurements in (C).

## DISCUSSION

In this study, we focused on the molecular basis underlying the strong temperature-sensitivity of Sup35NM droplet formation. Our results demonstrate that Tyr residues, which are abundant in the LS region of the N domain, play a crucial role in conferring this temperature-sensitivity. The data strongly suggest that Tyr-Tyr interactions stabilize both the local structure of the monomer and the droplet state. When these interactions become excessively strong, rapid droplet gelation occurs, leading to a reduction in temperature sensitivity. These findings suggest that the amino acid sequence of the Sup35 N domain is finely tuned to achieve an optimal interaction strength, determined by both the number and the spatial arrangement of Tyr residues, thereby enabling precise cellular regulation.

Although many intrinsically disordered proteins (IDPs) lack stable secondary structure in their monomeric state, they do not behave as ideal random coils. Instead, they adopt a sequence-dependent compact conformation (*42–44*). Importantly, the degree of compaction is closely linked to LLPS propensity, and LLPS-prone IDPs such as FUS-LCD and hnRNPA1/2-LCD are known to exhibit relatively compact conformations (*42, 44*). Previous studies have identified Tyr residues as one of the contributors to such compaction (*44, 45*). The LS region of Sup35NM, which is rich in Tyr and lacks defined secondary structure, resembles the compaction observed in other LLPS-associated IDPs. Notably, Tyr-mediated compaction is temperature-dependent, becoming more compact at low temperatures and more extended at higher temperatures (*44*). We propose that this temperature-responsive conformational behavior underlies the strong temperature-sensitivity of LLPS-prone IDPs. Sup35NM droplet formation exhibits sharp temperature-dependence, with rapid changes in droplet abundance over a narrow temperature range. At low temperatures, Tyr residues in the LS region promote chain compaction and form local clusters of densely packed Tyr residues. Interactions among these clusters facilitate droplet formation, and electrostatic interactions in the M domain further enhance droplet stability. As the temperature increases, Tyr clusters become destabilized due to chain expansion, while electrostatic interactions in the M domain are simultaneously weakened (*27*). Consequently, the Tyr-mediated interactions can no longer maintain the droplet state, leading to droplet dissociation (fig. S7). Interestingly, the isolated M domain can form droplets over a broad temperature range, whereas Sup35NM droplets dissociate at significantly lower temperatures (Fig. 1, A and B). This discrepancy may be explained by cation-π interactions between the 24 Lys residues in the M domain and the 20 Tyr residues in the N domain. Such N-M interactions may suppress M-M interactions, thereby enhancing overall temperature sensitivity (fig. S7). Supporting this model, mixing isolated N and M domains results in droplet formation containing both domains (fig. S8), suggesting direct interdomain interactions. Because temperature sensitivity is closely linked to cellular responsiveness to environmental stimuli, the strong temperature sensitivity of Sup35NM droplet formation may allow Sup35 to act as a sensitive sensor of environmental fluctuations.

Regarding the relationship between Tyr content and temperature-sensitivity, our previous characterization of Sup35NM variants from heterologous yeast strains also provides valuable insights (*36*). Sup35NM from budding yeast *Saccharomyces cerevisiae* (SC), *Kluyveromyces lactis* (KL), and *Candida albicans* (CA) exhibited strong temperature-sensitivity in droplet formation, whereas Sup35NM from the fission yeast *Schizosaccharomyces pombe* (SP) showed weak temperature-sensitivity, with a monotonic temperature dependence (*36*). Notably, the N domains of SC-, KL-, and CA-Sup35 contain 20 (16.2% of 123 residues), 24 (17.5% of 137 residues), and 18 (12.6% of 143 residues) Tyr residues, respectively, whereas SP-Sup35 contains only 6 Tyr residues (7.2% of 83 residues). These findings suggest that a threshold number of Tyr residues may be required to achieve sufficient IDP compaction, thereby conferring strong temperature-sensitivity to phase-separated droplets. Strikingly, the prion-like domains of the human FUS and hnRNPA1, which are linked to neurodegenerative diseases, contain 24 Tyr residues out of 163 residues (14.7%), and 20 Tyr/Phe residues (14.9%) out of 134 residues, respectively. These aromatic residue contents are nearly equivalent to those found in the N domain of budding yeast Sup35, suggesting that potential role for aromatic residues in these proteins in temperature sensitivity and response to environmental stimuli.

While this study has clarified the functional significance of Tyr residues, the roles of other amino acid residues remain largely unexplored. Prion-like domains are also enriched in small residues like glycine and serine, as well as polar residues such as asparagine and glutamine. These small residues are thought to enhance conformational flexibility. Although the specific contributions of polar residues to droplet formation remain unclear, their abundance and evolutionary conservation suggest that these residues may participate in fine-tuning phase behavior. Elucidating how these additional residues cooperate with aromatic interactions will be an important direction for future studies.

The Sup35 amyloid is a determinant of heritable [*PSI*^*+*^] strains, and no amyloid-associated cytotoxicity has been reported. Compared with amyloid-free [*psi*^*−*^] strains, [*PSI*^*+*^] strains have been reported to exhibit a survival advantage under stressful conditions (*46–48*). Thus, intracellular Sup35 amyloid formation could be beneficial as a cellular survival strategy under stress. Phase-separated droplet formation of Sup35 has also been observed under stress conditions (*17*) and is likely to contribute to stress tolerance. Although intermolecular interactions among the N domain are essential for both amyloid and droplet formation, these two assemblies involve distinct regions: droplet formation is primarily driven by the LS region located in the central part of the N domain, whereas amyloid formation is governed by the terminal regions, AM1 and AM2 (*30*). Such a division of roles may allow a pathway linking droplets to amyloids. Indeed, it has been reported that conversion from [*psi*^*−*^] to [*PSI*^*+*^] increases under various stress conditions (*49, 50*), which could reflect a relationship between droplet formation and amyloid emergence. Taken together, these observations suggest a potential model in which yeast cells initially form Sup35 droplets in response to acute stress and dissolve them upon stress relief, whereas under prolonged stress, droplets may mature into amyloids, thereby enabling a switch to a long-term adaptive state.

## MATERIALS AND METHODS

### Protein Expression and Purification

Wild-type Sup35NM and its mutants were cloned into the pET29b vector with a C-terminal His7-tag and expressed in *E. coli* BL21(DE3) cells. Lysates were clarified by centrifugation and applied to Ni-NTA agarose. Proteins were eluted with 250 mM imidazole under denaturing conditions. Further purification was performed by ion exchange chromatography under denaturing conditions. Sup35NM and Sup35M were purified via HiTrap SP (cation exchange), while Sup35N was recovered from the flow-through of HiTrap Q (anion exchange). Purified proteins were concentrated in 6 M guanidine hydrochloride.

### Sample Preparation for Droplet Formation Assays

To prepare samples for droplet formation, highly concentrated protein stock solutions were diluted with 20 mM sodium phosphate buffer (pH 7.0). The diluted samples were centrifuged at 21,000 × g for 30 minutes at room temperature to remove pre-formed aggregates. The final reaction mixture was adjusted to contain 10 μM protein, 10 mM sodium phosphate buffer (pH 7.0), and 10–20% (w/v) polyethylene glycol (PEG, average molecular weight 20,000, Nacalai Tesque). All sample mixing steps were performed on a 70 °C heat block to minimize premature phase separation. After mixing, the samples were cooled to room temperature and transferred into tubes or 96-well plates for subsequent analysis.

### Turbidity Measurement

All protein samples were prepared at 70 °C to minimize premature phase separation. Samples were dispensed into 96-well plates, cooled to 18 °C, sealed, and then placed in a microplate reader (EPOCH2, BioTek) pre-equilibrated at 20 °C. Turbidity was monitored by measuring absorbance at 500 nm. Measurements were performed over a temperature range of 20 °C to 60 °C in 5 °C increments. At each temperature, samples were incubated for 20 min prior to measurement to ensure thermal equilibration. For conditions containing additives such as NaCl or urea, which enhance aggregation propensity, measurements were conducted in two separate temperature ranges (e.g., 20–45 °C and 35–60 °C), and the datasets were combined to generate the full temperature profile. Each sample was measured in technical duplicates. All experiments were independently repeated three times (n = 3), and data are presented as mean ± s.e.m.

### Blue-native polyacrylamide gel electrophoresis

Blue native polyacrylamide gel electrophoresis (Blue-Native-PAGE) was performed using 10– 20% gradient gels without sodium dodecyl sulfate (SDS) (ATTO). Protein samples in 50% glycerol were thoroughly mixed with a native sample buffer containing 5% Coomassie Brilliant Blue G-250 (CBB G-250) and 100 mM Bis-Tris (pH 7.0), and then loaded onto the gel.

Electrophoresis was carried out using a cathode buffer consisting of 50 mM tricine, 15 mM Bis-Tris, and 0.02% CBB G-250 (pH 7.0), and an anode buffer consisting of 50 mM Bis-Tris (pH 7.0). The gel was run at a constant voltage of 100 V for 1 hour, followed by 150 V until the dye front reached the bottom of the gel. After electrophoresis, proteins were visualized by staining the gel with staining buffer (Nacalai Tesque).

### Optical and fluorescence microscopy

Optical microscopy was performed using a Eclipse LV 100POL (Nikon) microscope in bright-field mode. Observations were conducted using either a 40× or 100× objective lens. The microscope stage was equipped with a Peltier-based temperature control system, enabling precise regulation of the sample temperature during imaging.

Fluorescence imaging was performed using a BZ-X710 (KEYENCE) fluorescence microscope, equipped with a 100× objective lens. Samples were stained with Thioflavin T (ThT). GFP filter sets were used for excitation and emission to detect the respective fluorophores. Droplets images were acquired under identical exposure settings to allow comparison between samples.

### Amyloid formation kinetics measurements

Amyloid formation kinetics were monitored using thioflavin T (ThT) fluorescence. Protein samples (5 μM) were prepared in buffer containing 10 mM sodium phosphate (pH 7.0), 20% (w/v) polyethylene glycol (PEG 20,000), and 2.5 μM ThT. All samples were prepared on a 70 °C heat block to minimize premature phase separation or aggregation. Samples were dispensed into 348-well plates, cooled to room temperature, and placed in a fluorescence microplate reader pre-equilibrated at 24 °C. Fluorescence was measured with an excitation wavelength of 445 nm and an emission wavelength of 485 nm. Measurements were recorded every 30 min throughout the incubation period. Each condition was measured in technical triplicate. All experiments were independently repeated three times (n = 3), and data are presented as mean ± s.e.m.

### Fluorescence recovery after photobleaching (FRAP)

FRAP experiments were conducted using droplets that had been incubated for more than one hour after formation. Samples were prepared on glass-bottom dishes and imaged using a confocal laser scanning microscope (LSM 710, Carl Zeiss AG, Jena, Germany). Alexa Fluor 488-labeled proteins were used for fluorescence observation. A circular region within a droplet was photobleached using a 488 nm and 440 nm laser. Fluorescence recovery was monitored for a total duration of 80 seconds. All experiments were independently repeated seven times (n = 7), and data are presented as mean ± S.D.

### Circular dichroism (CD) measurement

CD spectra were recorded using a J-1100 CD spectropolarimeter (JASCO) equipped with a Peltier temperature controller. Protein samples were prepared in 10 mM phosphate buffer (pH 7.0) at a final concentration of 10 µM. Spectra were recorded at 20°C and 60°C using a quartz cuvette with a 1 mm path length.

## Funding

MEXT/JSPS KAKENHI 20J40038 (Y.O.)

MEXT/JSPS KAKENHI 20K06525 (Y.O.)

MEXT/JSPS KAKENHI 24K09363 (Y.O.)

MEXT/JSPS KAKENHI 20J21732 (S.N.)

MEXT/JSPS KAKENHI 22K18316 (M.F.)

MEXT/JSPS KAKENHI 20H03224 (E.C.)

MEXT/JSPS KAKENHI 25K02243 (E.C.)

Naito Foundation (Y.O.)

## Author contributions

Conceptualization: Y.O.

Methodology: Y.O., E.C., K.S., H.T.

Investigation: Y.O., S.N., Y.M.

Analyze and Discussion: Y.O., S.N., Y.M., M.F., K.S., E.C., H.T.

Writing: Y.O., H.T.

## REFERENCES

1. T. Ura, A. Kagawa, N. Sakakibara, H. Yagi, N. Tochio, T. Kigawa, K. Shiraki, T. Mikawa, Activation of L-lactate oxidase by the formation of enzyme assemblies through liquid–liquid phase separation. Sci. Rep. 13, 1435 (2023).

2. B. G. O’Flynn, T. Mittag, The role of liquid–liquid phase separation in regulating enzyme activity. Curr. Opin. Cell Biol. 69, 70–79 (2021).

3. S. F. Banani, H. O. Lee, A. A. Hyman, M. K. Rosen, Biomolecular condensates: organizers of cellular biochemistry. Nat. Rev. Mol. Cell Biol. 18, 285–298 (2017).

4. S. Alberti, A. A. Hyman, Biomolecular condensates at the nexus of cellular stress, protein aggregation disease and aging. Nat. Rev. Mol. Cell Biol. 22, 196–213 (2021).

5. M. Kato, T. W. Han, S. Xie, K. Shi, X. Du, L. C. Wu, H. Mirzaei, E. J. Goldsmith, J. Longgood, J. Pei, N. V. Grishin, D. E. Frantz, J. W. Schneider, S. Chen, L. Li, M. R. Sawaya, D. Eisenberg, R. Tycko, S. L. McKnight, Cell-free formation of RNA granules: low complexity sequence domains form dynamic fibers within hydrogels. Cell 149, 753–67 (2012).

6. A. C. Murthy, G. L. Dignon, Y. Kan, G. H. Zerze, S. H. Parekh, J. Mittal, N. L. Fawzi, Molecular interactions underlying liquid−liquid phase separation of the FUS low-complexity domain. Nat. Struct. Mol. Biol. 26, 637–648 (2019).

7. W. Zheng, G. L. Dignon, N. Jovic, X. Xu, R. M. Regy, N. L. Fawzi, Y. C. Kim, R. B. Best, J. Mittal, Molecular details of protein condensates probed by microsecond long atomistic simulations. J. Phys. Chem. B 124, 11671–11679 (2020).

8. T. R. Peskett, F. Rau, J. O’Driscoll, R. Patani, A. R. Lowe, H. R. Saibil, A liquid to solid phase transition underlying pathological huntingtin exon1 aggregation. Mol. Cell 70, 588-601.e6 (2018).

9. S. Ray, N. Singh, R. Kumar, K. Patel, S. Pandey, D. Datta, J. Mahato, R. Panigrahi, A. Navalkar, S. Mehra, L. Gadhe, D. Chatterjee, A. S. Sawner, S. Maiti, S. Bhatia, J. A. Gerez, A. Chowdhury, A. Kumar, R. Padinhateeri, R. Riek, G. Krishnamoorthy, S. K. Maji, α-Synuclein aggregation nucleates through liquid–liquid phase separation. Nat. Chem. 12, 705–716 (2020).

10. P. Chakraborty, M. Zweckstetter, Role of aberrant phase separation in pathological protein aggregation. Curr. Opin. Struct. Biol. 82, 102678 (2023).

11. B. S. Visser, W. P. Lipiński, E. Spruijt, The role of biomolecular condensates in protein aggregation. Nat. Rev. Chem. 8, 686–700 (2024).

12. S. Qamar, G. Wang, S. J. Randle, F. Ruggeri, J. A. Varela, J. Lin, E. C. Phillips, A. Miyashita, D. Williams, F. Ströhl, W. Meadows, R. Ferry, V. J. Dardov, G. G. Tartaglia, L. A. Farrer, G. S. Schierle, C. F. Kaminski, C. E. Holt, P. E. Fraser, G. Schmitt-Ulms, D. Klenerman, T. Knowles, M. Vendruscolo, P. George-Hyslop, FUS phase separation is modulated by a molecular chaperone and methylation of arginine cation-π interactions. Cell 173, 720-734.e15 (2018).

13. X. Su, J. A. Ditlev, E. Hui, W. Xing, S. Banjade, J. Okrut, D. S. King, J. Taunton, M. K. Rosen, R. D. Vale, Phase separation of signaling molecules promotes T cell receptor signal transduction. Science 352, 595–599 (2016).

14. Y. Liu, W. Feng, Y. Wang, B. Wu, Crosstalk between protein post-translational modifications and phase separation. Cell Commun. Signal. 22, 110 (2024).

15. Z. Liu, S. Zhang, J. Gu, Y. Tong, Y. Li, X. Gui, H. Long, C. Wang, C. Zhao, J. Lu, L. He, Y. Li, Z. Liu, D. Li, C. Liu, Hsp27 chaperones FUS phase separation under the modulation of stress-induced phosphorylation. Nat. Struct. Mol. Biol. 27, 363–372 (2020).

16. F. Aryan, D. Detrés, C. C. Luo, S. X. Kim, A. N. Shah, M. Bartusel, R. A. Flynn, E. Calo, Nucleolus activity-dependent recruitment and biomolecular condensation by pH sensing. Mol. Cell 83, 4413-4423.e10 (2023).

17. T. M. Franzmann, M. Jahnel, A. Pozniakovsky, J. Mahamid, A. S. Holehouse, E. Nüske, D. Richter, W. Baumeister, S. W. Grill, R. V. Pappu, A. A. Hyman, S. Alberti, Phase separation of a yeast prion protein promotes cellular fitness. Science 359 (2018).

18. S. Cinar, H. Cinar, H. S. Chan, R. Winter, Pressure-sensitive and osmolyte-modulated liquid– liquid phase separation of eye-lens γ-crystallins. J. Am. Chem. Soc. 141, 7347–7354 (2019).

19. Z. Fetahaj, L. Ostermeier, H. Cinar, R. Oliva, R. Winter, Biomolecular condensates under extreme martian salt conditions. J. Am. Chem. Soc. 143, 5247–5259 (2021).

20. Y. Lin, D. S. Protter, M. K. Rosen, R. Parker, Formation and maturation of phase-separated liquid droplets by RNA-binding proteins. Mol. Cell 60, 208–19 (2015).

21. S. K. Rai, R. Khanna, A. Avni, S. Mukhopadhyay, Heterotypic electrostatic interactions control complex phase separation of tau and prion into multiphasic condensates and coaggregates. Proc. Natl. Acad. Sci. 120, e2216338120 (2023).

22. R. Sternke-Hoffmann, X. Sun, A. Menzel, M. D. S. Pinto, U. Venclovaite, M. Wördehoff, W. Hoyer, W. Zheng, J. Luo, Phase separation and aggregation of α-synuclein diverge at different salt conditions. Adv. Sci., e2308279 (2024).

23. A. S. Sawner, S. Ray, P. Yadav, S. Mukherjee, R. Panigrahi, M. Poudyal, K. Patel, D. Ghosh, E. Kummerant, A. Kumar, R. Riek, S. K. Maji, Modulating α-synuclein liquid–liquid phase separation. Biochemistry-us 60, 3676–3696 (2021).

24. H. Cinar, Z. Fetahaj, S. Cinar, R. M. Vernon, H. S. Chan, R. H. A. Winter, Temperature, hydrostatic pressure, and osmolyte effects on liquid–liquid phase separation in protein condensates: physical chemistry and biological implications. Chem. A Eur. J. 25, 13049–13069 (2019).

25. K. M. Ruff, S. Roberts, A. Chilkoti, R. V. Pappu, Advances in understanding stimulus-responsive phase behavior of intrinsically disordered protein polymers. J. Mol. Biol. 430, 4619–4635 (2018).

26. F. G. Quiroz, A. Chilkoti, Sequence heuristics to encode phase behaviour in intrinsically disordered protein polymers. Nat. Materials 14, 1164–1171 (2015).

27. D. J. Andlinger, U. Kulozik, Protein–protein interactions explain the temperature-dependent viscoelastic changes occurring in colloidal protein gels. Soft Matter 19, 1144–1151 (2022).

28. A. V. Grizel, N. A. Gorsheneva, J. B. Stevenson, J. Pflaum, F. Wilfling, A. A. Rubel, Y. O. Chernoff, Osmotic stress induces formation of both liquid condensates and amyloids by a yeast prion domain. J. Biol. Chem., 107766 (2024).

29. B. Grimes, W. Jacob, A. R. Liberman, N. Kim, X. Zhao, D. C. Masison, L. E. Greene, The properties and domain requirements for phase separation of the Sup35 prion protein in vivo. Biomolecules 13, 1370 (2023).

30. Y. Ohhashi, Y. Yamaguchi, H. Kurahashi, Y. O. Kamatari, S. Sugiyama, B. Uluca, T. Piechatzek, Y. Komi, T. Shida, H. Müller, S. Hanashima, H. Heise, K. Kuwata, M. Tanaka, Molecular basis for diversification of yeast prion strain conformation. Proc. Natl. Acad. Sci. USA 115, 2389–2394 (2018).

31. H. Konno, T. Watanabe-Nakayama, T. Uchihashi, M. Okuda, L. Zhu, N. Kodera, Y. Kikuchi, T. Ando, H. Taguchi, Dynamics of oligomer and amyloid fibril formation by yeast prion Sup35 observed by high-speed atomic force microscopy. Proc. Natl. Acad. Sci. 117, 7831–7836 (2020).

32. B. H. Toyama, M. J. S. Kelly, J. D. Gross, J. S. Weissman, The structural basis of yeast prion strain variants. Nature 449, 233–237 (2007).

33. O. D. King, A. D. Gitler, J. Shorter, The tip of the iceberg: RNA-binding proteins with prion-like domains in neurodegenerative disease. Brain Res. 1462, 61–80 (2012).

34. E. Chatani, K. Yuzu, Y. Ohhashi, Y. Goto, Current understanding of the structure, stability and dynamic properties of amyloid fibrils. Int. J. Mol. Sci. 22, 4349 (2021).

35. M. Fukuyama, S. Nishinami, Y. Maruyama, T. Ozawa, S. Tomita, Y. Ohhashi, M. Kasuya, M. Gen, E. Chatani, K. Shiraki, A. Hibara, Detection of fibril nucleation in micrometer-sized protein condensates and suppression of Sup35NM fibril nucleation by liquid–liquid phase separation. Anal. Chem. 95, 9855–9862 (2023).

36. Y. Ohhashi, S. Nishinami, K. Shiraki, E. Chatani, H. Taguchi, Low-complexity domains in phase-separated droplets suppress the amyloid formation of yeast prion Sup35. npj Biosensing 2, 13 (2025).

37. Y. Ohhashi, K. Ito, B. H. Toyama, J. S. Weissman, M. Tanaka, Differences in prion strain conformations result from non-native interactions in a nucleus. Nat. Chem. Biol. 6, 225–230 (2010).

38. K. Kamagata, M. Ariefai, H. Takahashi, A. Hando, D. R. G. Subekti, K. Ikeda, A. Hirano, T. Kameda, Rational peptide design for regulating liquid–liquid phase separation on the basis of residue–residue contact energy. Sci. Rep. 12, 13718 (2022).

39. A. Thomas, R. Meurisse, R. Brasseur, Aromatic side-chain interactions in proteins. II. Near-and far-sequence Phe-X pairs. Proteins 48, 635–644 (2002).

40. X. Gui, F. Luo, Y. Li, H. Zhou, Z. Qin, Z. Liu, J. Gu, M. Xie, K. Zhao, B. Dai, W. S. Shin, J. He, L. He, L. Jiang, M. Zhao, B. Sun, X. Li, C. Liu, D. Li, Structural basis for reversible amyloids of hnRNPA1 elucidates their role in stress granule assembly. Nat. Commun. 10, 2006 (2019).

41. I. Peran, T. Mittag, Molecular structure in biomolecular condensates. Curr. Opin. Struct. Biol. 60, 17–26 (2020).

42. G. Tesei, A. I. Trolle, N. Jonsson, J. Betz, F. E. Knudsen, F. Pesce, K. E. Johansson, K. Lindorff-Larsen, Conformational ensembles of the human intrinsically disordered proteome. Nature 626, 897–904 (2024).

43. J. A. Marsh, J. D. Forman-Kay, Sequence determinants of compaction in Intrinsically disordered proteins. Biophys. J. 98, 2383–2390 (2010).

44. E. W. Martin, A. S. Holehouse, I. Peran, M. Farag, J. J. Incicco, A. Bremer, C. R. Grace, A. Soranno, R. V. Pappu, T. Mittag, Valence and patterning of aromatic residues determine the phase behavior of prion-like domains. Science 367, 694–699 (2020).

45. K. Linhartova, F. L. Falginella, M. Matl, M. Sebesta, R. Vácha, R. Stefl, Sequence and structural determinants of RNAPII CTD phase-separation and phosphorylation by CDK7. Nat. Commun. 15, 9163 (2024).

46. S. S. Eaglestone, B. S. Cox, M. F. Tuite, Translation termination efficiency can be regulated in Saccharomyces cerevisiae by environmental stress through a prion-mediated mechanism. EMBO J. 18, 1974–1981 (1999).

47. H. L. True, S. L. Lindquist, A yeast prion provides a mechanism for genetic variation and phenotypic diversity. Nature 407, 477–483 (2000).

48. R. Halfmann, D. F. Jarosz, S. K. Jones, A. Chang, A. K. Lancaster, S. Lindquist, Prions are a common mechanism for phenotypic inheritance in wild yeasts. Nature 482, 363–368 (2012).

49. V. A. Doronina, G. L. Staniforth, S. H. Speldewinde, M. F. Tuite, C. M. Grant, Oxidative stress conditions increase the frequency of de novo formation of the yeast [PSI+] prion. Mol. Microbiology 96, 163–174 (2015).

50. J. Tyedmers, M. L. Madariaga, S. Lindquist, Prion switching in response to environmental stress. Plos Biol 6, e294 (2008).

