## Supplemental Figure 1 - 8 for "Local aromatic interactions define temperature sensitivity of phase separation in an intrinsically disordered protein"

**This PDF file includes:**

Figs. S1 to S8

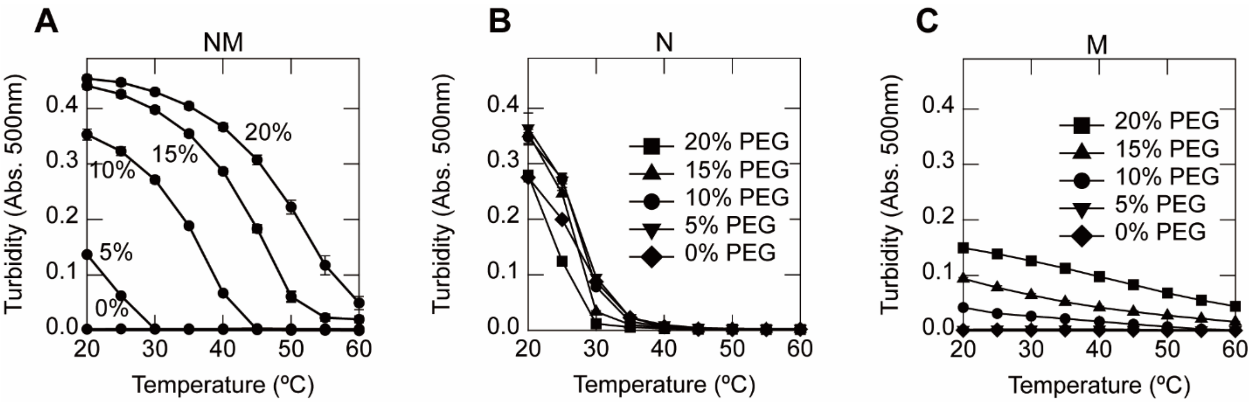


**Fig. S1. PEG 20,000 concentration dependence of droplet formation of Sup35NM (A), N (B), and M (C) domains measured by turbidity.**

| | | |
|---|---|---|
| AM1 | | MSDSNQGNNQ QNYQQYSQNG NQQQGNNRYQ GYQAYNAQAQ PAGGY |
| LS | (M) | AGGYYQNYQG YSGYQQGGYQ QYNPDAGYQQ QYNPQGGYQQ YNPQGGYQQ |
| AM2 | (M) | QQYNPQGGYQ QQFNPQGGRG NYKNFNYNNN LQGYQAGFQP QSQG |

**Fig. S2. Amino acid sequences of the three regions within the Sup35 N domain: AM1, LS, and AM2. Tyrosine residues are shown in red.**

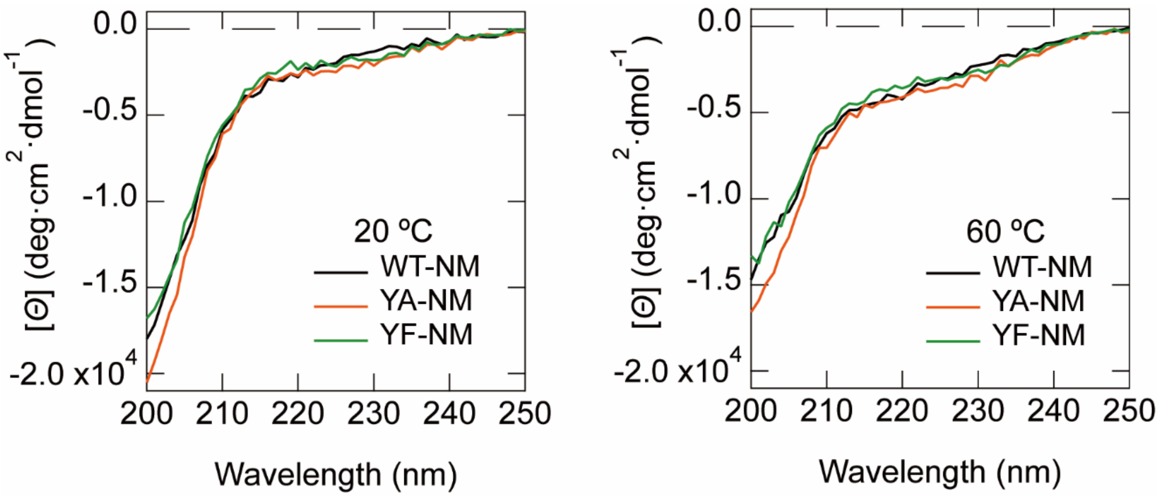


**Fig. S3. Far-UV circular dichroism spectra of WT-NM, YA-NM, and YF-NM at 20 °C (left) and 60 °C (right).**

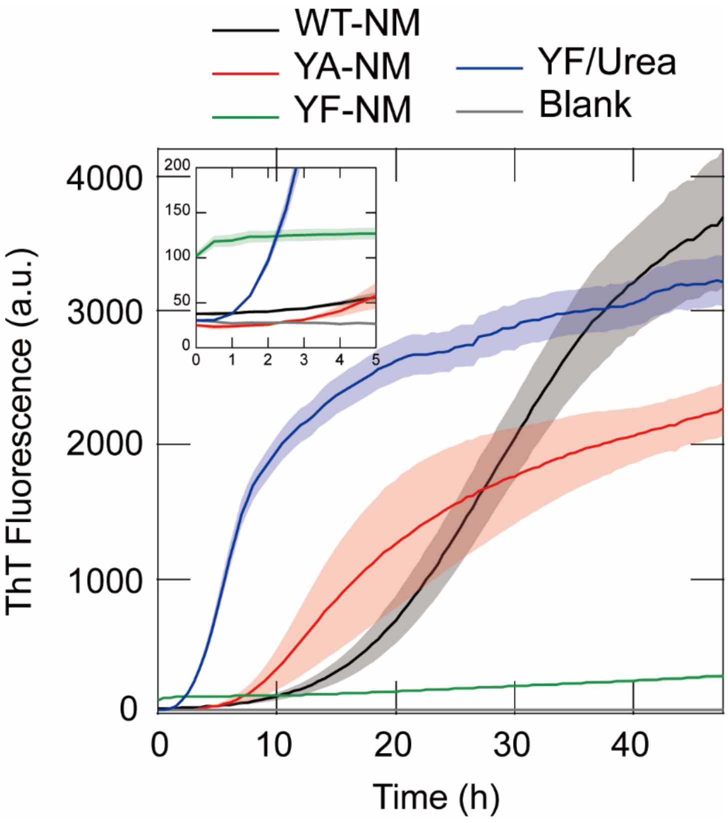


**Fig. S4. Time course of ThT fluorescence intensity for WT-NM, YA-NM, and YF-NM droplets in the absence or presence of 2 M urea, measured at 24 °C.**
Data represent mean ± s.e.m. (n = 3), with error shown as shaded regions. The inset shows a magnified view of the first 5 h.

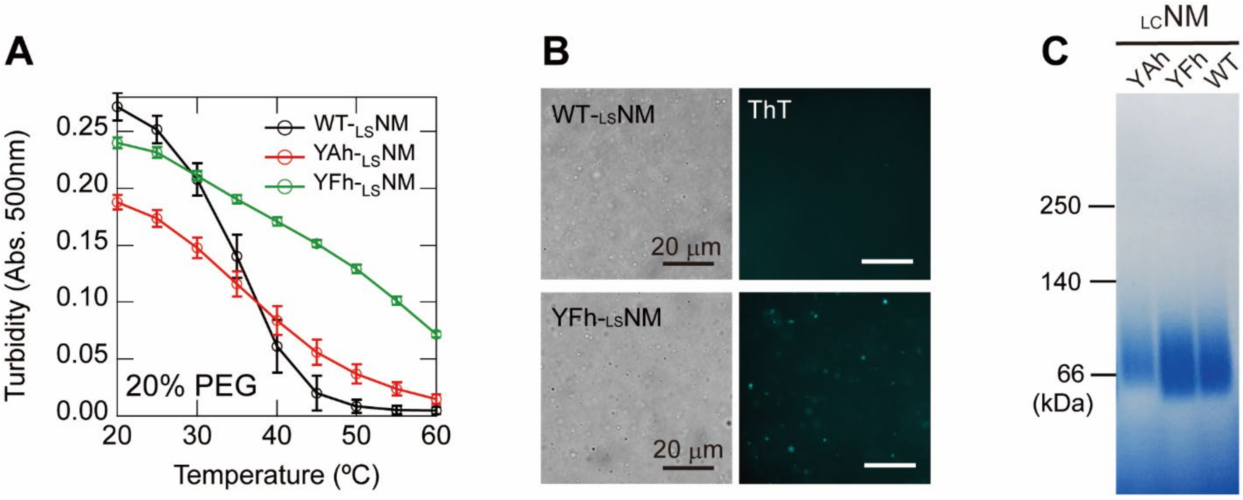


**Fig. S5. Properties of Ala- and Phe-substituted LSNM mutants reveal altered phase behavior.**

(**A**) Temperature-dependent turbidity profiles of WT-LSNM, YAh-LSNM, and YFh-LSNM in 20% (w/v) PEG 20,000.

(**B**) ThT fluorescence images of droplets formed by WT-LSNM and YFh-LSNM in 20% (w/v) PEG 20,000. Scale bars, 20 µm.

(**C**) Blue-Native PAGE analysis of WT-LSNM, YAh-LSNM, and YFh-LSNM.

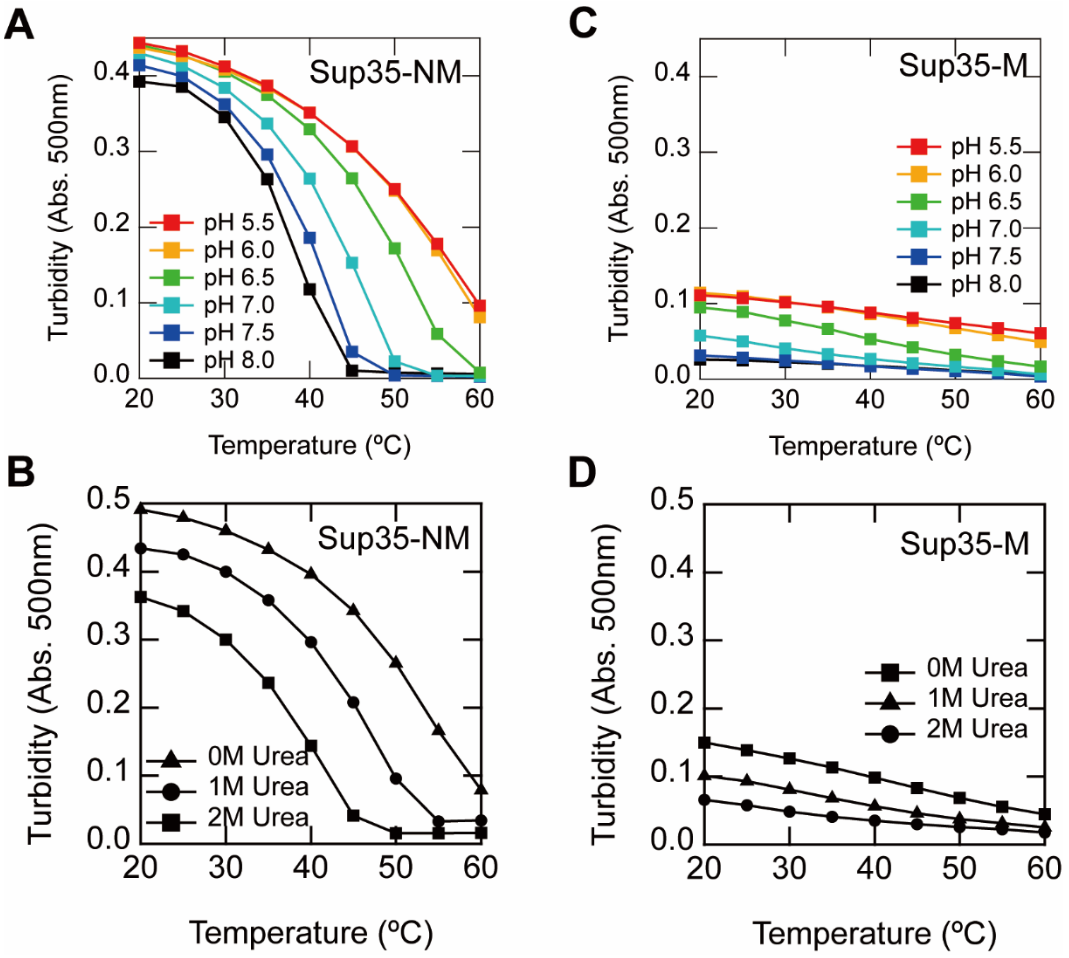


**Fig. S6. Temperature sensitivity cooperates with additional environmental factors.**
(**A**) pH dependence of Sup35NM droplet formation measured by turbidity.
(**B**) Urea concentration dependence of Sup35NM droplet formation measured by turbidity.
(**C**) pH dependence of Sup35M droplet formation measured by turbidity.
(**D**) Urea concentration dependence of Sup35M droplet formation measured by turbidity.

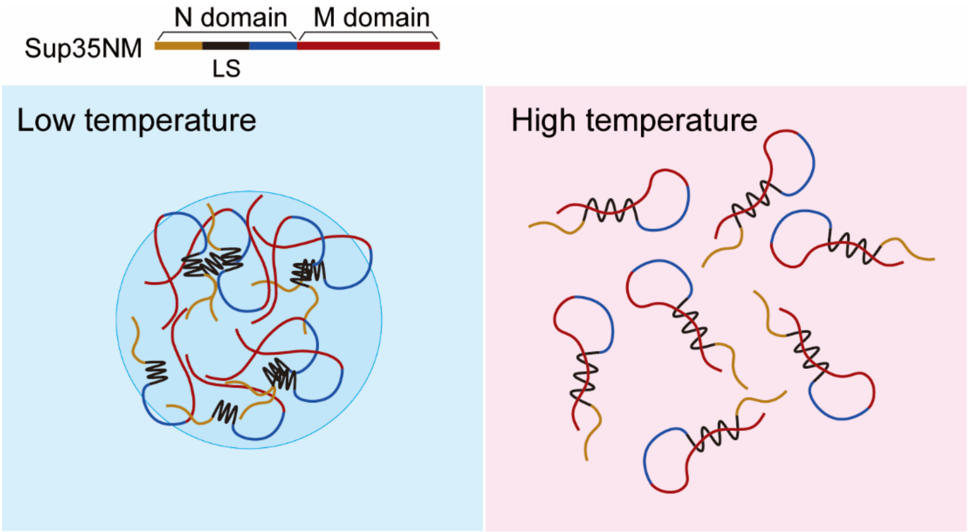


**Fig. S7. Temperature sensitivity cooperates with additional environmental factors.** At low temperature, the LS region of the N domain adopts compact local structures in which tyrosine residues cluster and mediate intermolecular interactions, while M domains interact electrostatically (left). At elevated temperatures, loosening of the local structures disrupts tyrosine clustering, leading to loss of intermolecular interactions. As N–N interactions weaken, the LS region engages in intramolecular cation–π interactions with the M domain, thereby disrupting M–M interactions.

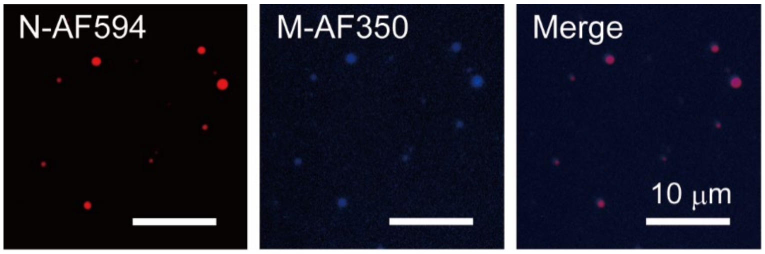


**Fig. S8. Representative fluorescence images of droplets formed from a mixture of Sup35N-Alexa Fluor 594 and Sup35M-Alexa Fluor 350, showing colocalization of both proteins within the droplets.**
Scale bars, 10 μm.
